# Improving Probabilistic Infectious Disease Forecasting Through Coherence

**DOI:** 10.1101/2019.12.27.889212

**Authors:** Graham Casey Gibson, Kelly R. Moran, Nicholas G. Reich, Dave Osthus

## Abstract

With an estimated $10.4 billion in medical costs and 31.4 million outpatient visits each year, influenza poses a serious burden of disease in the United States. To provide insights and advance warning into the spread of influenza, the U.S. Centers for Disease Control and Prevention (CDC) runs a challenge for forecasting weighted influenza-like illness (wILI) at the national and regional level. Many models produce independent forecasts for each geographical unit, ignoring the constraint that the national wILI is a weighted sum of regional wILI, where the weights correspond to the population size of the region. We propose a novel algorithm that transforms a set of independent forecast distributions to obey this constraint, which we refer to as probabilistically coherent. Enforcing probabilistic coherence led to an increase in forecast skill for 90% of the models we tested over multiple flu seasons, highlighting the importance of respecting the forecasting system’s geographical hierarchy.

**Author Summary:** Seasonal influenza causes a significant public health burden nationwide. Accurate influenza forecasting may help public health officials allocate resources and plan responses to emerging outbreaks. The U.S. Centers for Disease Control and Prevention (CDC) reports influenza data at multiple geographical units, including regionally and nationally, where the national data are by construction a weighted sum of the regional data. In an effort to improve influenza forecast accuracy across all models submitted to the CDC’s annual flu forecasting challenge, we examined the effect of imposing this geographical constraint on the set of independent forecasts, made publicly available by the CDC. We developed a novel method to transform forecast densities to obey the geographical constraint that respects the correlation structure between geographical units. This method showed consistent improvement across 90% of models and that held when stratified by targets and test seasons. Our method can be applied to other forecasting systems both within and outside an infectious disease context that have a geographical hierarchy.

## 1 Introduction

Seasonal influenza is a persistent and serious contributor to global morbidity and mortality, hospitalizing over half a million people in the world every year [1]. The United States alone reported approximately 80,000 influenza related mortalities in the 2017/2018 influenza season, with most serious consequences for vulnerable populations such as children or the elderly [2].

As part of a larger forecasting initiative, the U.S. Centers for Disease Control and Prevention (CDC) hosts an annual influenza forecasting challenge called the FluSight challenge open to the public [3][4]. As part of this challenge, forecasters supply probabilistic forecasts for short-term and seasonal targets at both the national and regional levels corresponding to weighted influenza-like illness (wILI), which measures the proportion of outpatient doctor visits at reporting health care facilities where the patient had an influenza-like illness (ILI), weighted by state population. At the national level, wILI can be directly computed using state population weighted ILI or it can be equivalently computed using regional population weighted wILI. The CDC estimates ILI as the ratio of patients presenting with a cough and fever equal to or above 100° Fahrenheit over the total number of patients presenting at health care providers [5]. Participants in the FluSight challenge have harnessed a variety of models and methods to forecast the targets under consideration, which include both short-term forecasts and seasonal targets. These efforts have included time series models [6], mechanistic disease transmission models [7][8], and machine learning techniques [9][10][11][12]. Teams have also incorporated external data, such as internet search queries or point of care data, to improve forecasts [13][14][15][16]. FluSight challenge participation has grown in popularity since the inaugural challenge in 2013, with twenty-four teams submitting forecasts from thirty-three models for the 2018/2019 season [17]. Model submission files from the past FluSight challenges are publicly available [17], providing the opportunity for retrospective analysis and the potential for improved forecasting.

Multiple procedures have been proposed in the literature to transform independently generated incoherent forecasts into coherent forecasts, also called forecast reconciliation [18] [19]. Projection matrix forecasting is a popular coherence forecasting approach [20]. This approach uses a matrix projection of the original set of forecasts onto a subspace that respects the known hierarchical relationship of the forecasting system. This approach uses forecasts for all levels of the hierarchy and does not discard any information as opposed to say a bottom-up approach, where the national forecast is ignored and the estimate is simply a linear combination of regional estimates. However, the demonstrated benefits of coherence in the point prediction setting do not necessarily translate to the probabilistic forecasting realm [19]. We propose a novel algorithm that transforms a set of independent forecast densities to satisfy the coherence property and demonstrate that the resulting collection of forecast densities has a consistently higher forecast skill when broken down by season and target.

### 1.1 US ILI Surveillance Data

For the FluSight challenge, the CDC provides wILI data at both the national level and broken down into 10 Health and Human Services (HHS) regions, mostly organized by geographic proximity. The data are reported on a weekly basis and extend from 1997 to the present. Example data for the 2016/2017, 2017/2018, and 2018/2019 seasons are shown in Figure 1. As noted in the Figure 1, wILI varies by region but maintains a relatively consistent winter peak. The CDC reports the wILI data using epidemic weeks, called epiweeks, instead of calendar weeks [21]. This allows for consistent week numbering across multiple seasons. Epiweek 40 is usually the first week of October, the start of the flu season, and epiweek 20 usually falls in May, marking the end of the season.

**Figure 1:**
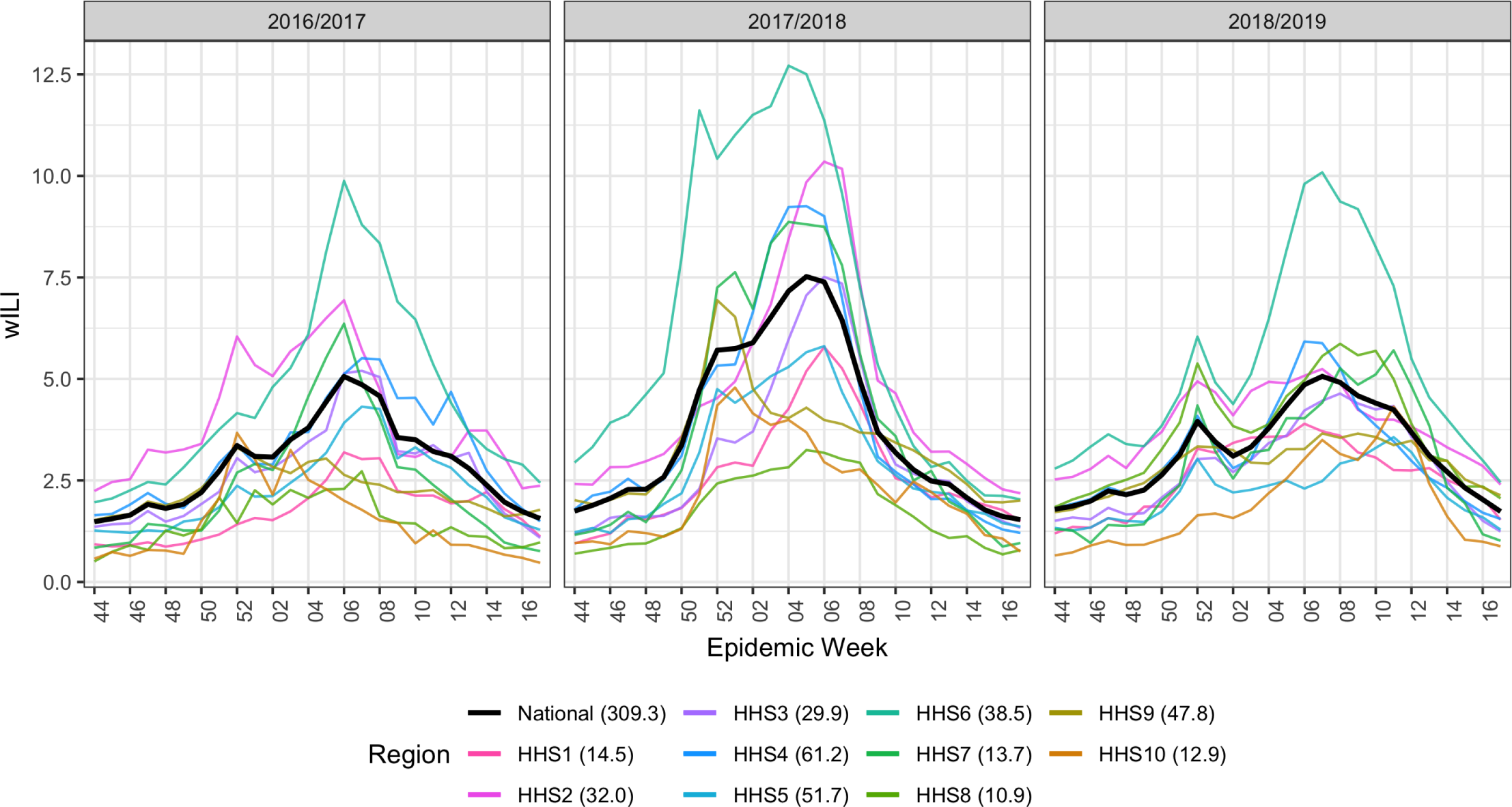
Data example for the three test seasons under consideration (2016/2017, 2017/2018, 2018/2019) season for all 10 Health and Human Services (HHS) regions and the national level. At any given epiweek, the national wILI (black) is a weighted sum of regional wILI, where the weights correspond to the population size of the region. We can see that wILI is highly seasonal and varies heavily by region. Region population sizes (in millions) are given next to the region in the legend.

### 1.2 Pointwise Forecast Coherence

The partitioning of national data into HHS regions facilitates geographically localized forecasts, augmenting their usability to local public health officials. A consequence of this partitioning, however, is the creation of a hierarchical structure in the forecasting system. Namely, national wILI data is a linear combination of HHS regional wILI data. Region population sizes (in millions) are given next to the region in the legend of Figure 1 as reported by the 2010 U.S. Census [22].

We notate the true wILI value as *y*_*r,s,w*_ ∈ [0, 100], a percentage for region *r* in flu season *s* corresponding to epiweek *w*. Throughout the paper, *r* = 11 corresponds to the nation, while *r* = 1, 2, …, 10 corresponds to HHS region *r*. Let *α*_*r*_ ∈ [0, 1] be a weight corresponding to HHS region *r*, proportional to the population of HHS region *r*, such that 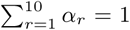. The hierarchical nature of the national/regional partitioning of forecasts for any season and epiweek is equivalent to the following constraint:

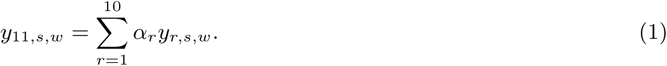

For convenience, define the collection of point forecasts for all regions for season *s* and epiweek *w* as

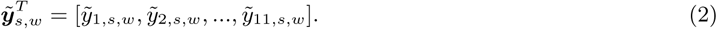

We say that the forecast 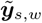 is *coherent* if

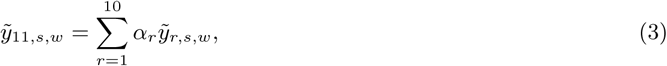

and 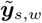 is *incoherent* if

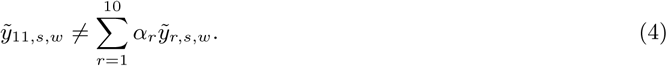

Though the existence of the hierarchical structure in the wILI forecasting system is know, many FluSight challenge forecasts are made independently. Forecasting at geographic regions independently provides the forecaster more flexibility to cater models to specific regions or avoid modeling correlation between regions explicitly, but leaves the resulting forecasts vulnerable to incoherence as the true coherent data generating process is not respected. In this paper, we use 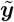 to represent independently generated and (likely) incoherent forecasts and ***ŷ*** to represent coherent forecasts.

For an independently generated set of forecasts 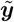, the corresponding coherent projection matrix forecast ***ŷ*** is

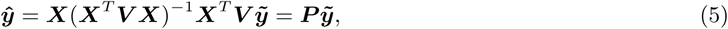

where ***X*** is a design matrix corresponding to the hierarchical relationship of the data generating process and ***V*** is a weight matrix. Specifically for the FluSight challenge,

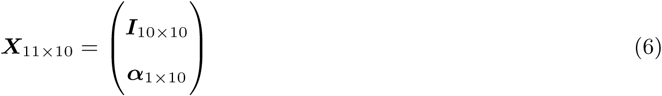

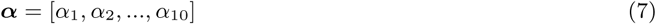

where *α*_*r*_ is the weight for the *r*^*th*^ region.

A special case of projection matrix forecasting is when the ordinary least squares (OLS) projection matrix is used, produced by setting ***V*** equal to the identity matrix ***I*** in Equation 5. This special case has the property that the resulting coherent point forecast ***ŷ*** has mean squared error (MSE) no worse than the independently generated forecast 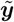 [19]. That is, when ***V*** = ***I*** in Equation 5,

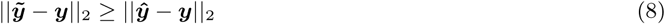

for any ***y*** and 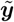 (See Appendix for proof).

For illustration and clarity, consider an example with two low level regions (HHS1 and HHS2) and one top level region (Nation). This example is illustrated in Figure 2.

**Figure 2:**
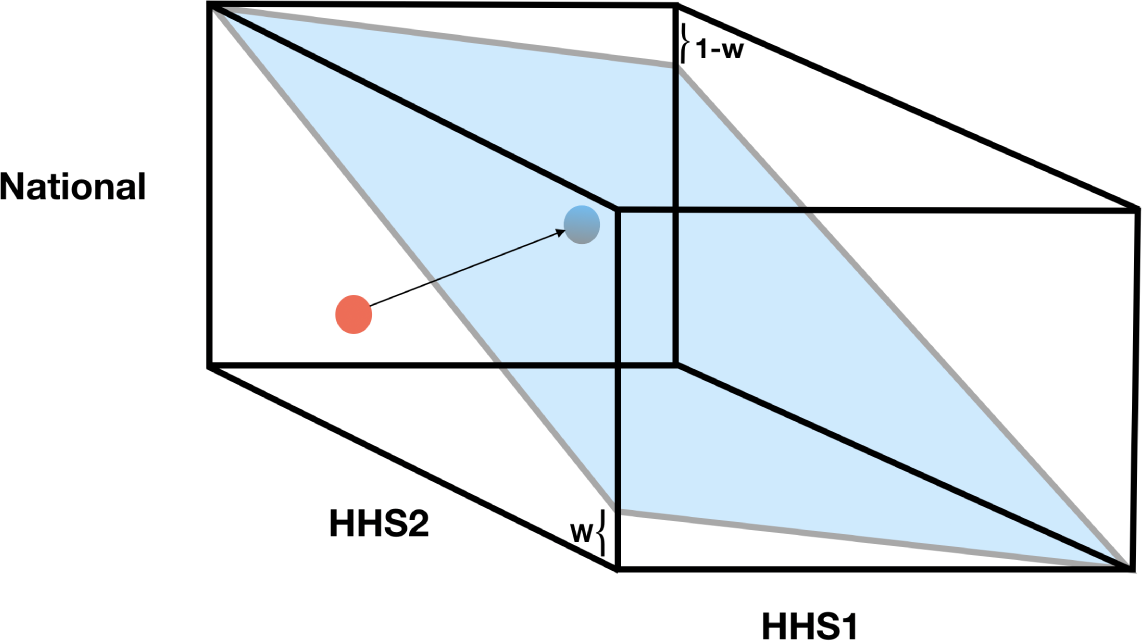
Projection of original forecasts (red) onto space satisfying constraint of regional level forecasts summing to national level (blue). The three different projection methods defined by the various choices of weight matrices result in different projections of the red point onto the blue plane. The blue plane represents the set of points that satisfy the coherence constraint, namely that the weighted combination of region-level forecasts equals the National level forecast. In particular, the ordinary least squares projection (OLS) projects the red point onto the blue point such that the resulting distance is minimized, leading to an orthogonal projection.

Assume

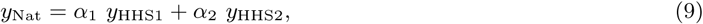

where *α*_1_ = *α*_2_ = 0.5. Let the true values be

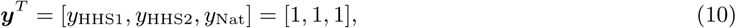

and assume the independently generated forecasts are

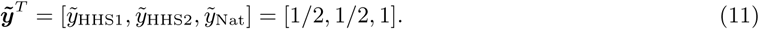

Notice that the independently generated forecasts are incoherent, as

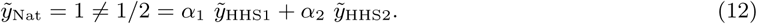

The MSE for 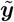 is

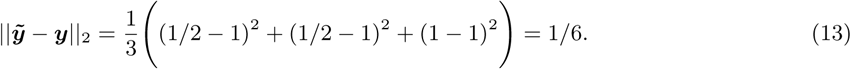

The OLS projection matrix forecast is ***ŷ*** = [2/3, 2/3, 2/3], computed as

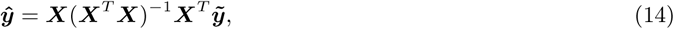

where

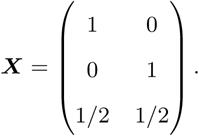

The projection matrix forecast ***ŷ*** is, by construction, coherent. The effect of the projection matrix forecast is a reduction in MSE over 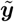, where the MSE for ***ŷ*** is

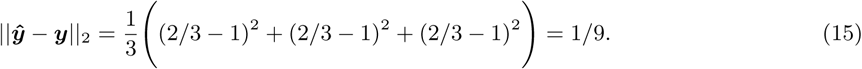

Note the MSE for *ŷ*_HHS1_ and *ŷ*_HHS2_ improved relative to 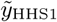 and 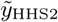, respectively, while the MSE for *ŷ*_Nat_ got worse relative to 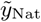, resulting in an overall, but not uniform, improvement in MSE for ***ŷ*** relative to 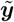. The projection matrix forecast is a useful tool for transforming independently generated, incoherent forecasts into coherent forecasts. When ***V*** = ***I*** in Equation 5, the MSE of the resulting ***ŷ*** is guaranteed to be no greater than the MSE of 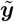. When ***V*** ≠ ***I*** in Equation 5, the resulting forecast is still coherent, but no such guarantee of MSE improvement exists.

### 1.3 The FluSight Forecast Coherence Dilemma

Unlike the situations demonstrated in the point forecasting literature, forecasts for the FluSight challenge are required to be probabilistic, not point estimates, and the probabilistic forecasts are evaluated using a multi-bin scoring rule, not MSE. In practice, probabilistic forecasts are generated as a collection of *n* forecast samples for *y*_*r,s,w*_. In this paper, we use the index *i* = 1, 2, …, *n* to denote draw *i* from the forecast distribution, resulting in a collection of realizations 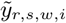. In probabilistic settings we can no longer rely on the coherence definition given in Equation 3. Although various definitions for probabilistic coherence exist [23][18], in this paper, we choose the intuitive presentation of Gamakumara et. al [23]. We say that the density 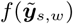 is *probabilistically coherent* if

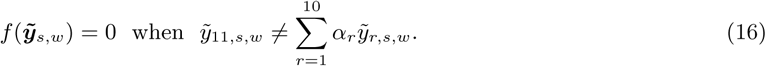

Here *f* represents the joint density over all regions and the national level of the hierarchy. Note the close correspondence with the point forecast definition. However, probabilistic coherence does require specification of a joint distribution over regions. Intuitively, this definition says that any point in the support of *f* that does not obey point-wise coherence is assigned zero probability, (i.e., has measure zero).

We score the probabilistic forecasts using multi-bin skill, rather than multi-bin log score as used in the FluSight challenge. Multi-bin skill is on the interpretable [0,1] probability scale, unlike the multi-bin log score. Like log score, skill is a thresholding scoring rule, where a forecast is deemed correct if it is within a certain distance of the truth. The probability assigned to the true target *Z*_*t*_ (e.g., a one-week-ahead forecast) corresponding to region *r* in flu season *s* and epiweek *w* is computed as

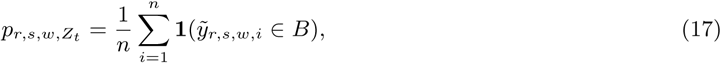

where *B* = [*Z*_*t*_ − *b*, *Z*_*t*_ + *b*] for the true target *Z*_*t*_ and a predefined threshold *b*. For short-term forecasts (e.g., one through four-week-ahead targets), *b* is set equal to 0.5. Using a thresholding evaluation metric breaks the guaranteed equal or improved performance of the coherent forecasts when evaluated with MSE.

To see why, consider again the example from Section 1.2 with ***y***^*T*^ = [1, 1, 1]. For *i* = 1, 2, …, *n* = 10000, we independently draw

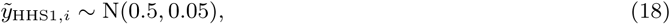

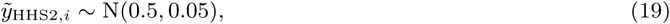

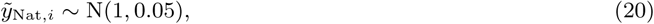

defining 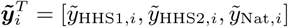 and 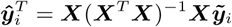. Figure 3 shows the *n* draws of 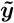 and ***ŷ***. The skill counts all forecasts within a predefined threshold of the true value as correct and all forecasts outside the predefined threshold as incorrect. For this example, the correct forecasts must fall within plus or minus 0.1 of the true value ***y***. For every realization *i*, the MSE for ***ŷ***_*i*_ is less than the MSE of 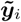. However, the skill for the coherent forecasts is 0 while the skill for incoherent forecasts is 0.32 and the skill for the incoherent forecasts is always better than or equal to the skill for the coherent forecasts. This is because all of the correct forecasts nationally are projected out of the correct region while the incorrect regional forecasts are moved closer to the correct region, but not close enough to fall inside it. The result is a collection of coherent forecasts with better (lower) MSE, but also lower forecast skill than the incoherent forecasts.

### 1.4 Problem Statement

On one hand, we have a guarantee that the MSE of point forecasts projected into the data generating process space can get no worse under the OLS projection method. On the other, we have an explicit example of forecast skill decreasing as a result of forecasts projected into the data generating process space. This seeming inconsistency leads us to the central question of this paper: Can probabilistic forecast coherence be used to improve forecast skill when forecasting influenza? The remainder of this paper will investigate this question.

**Figure 3:**
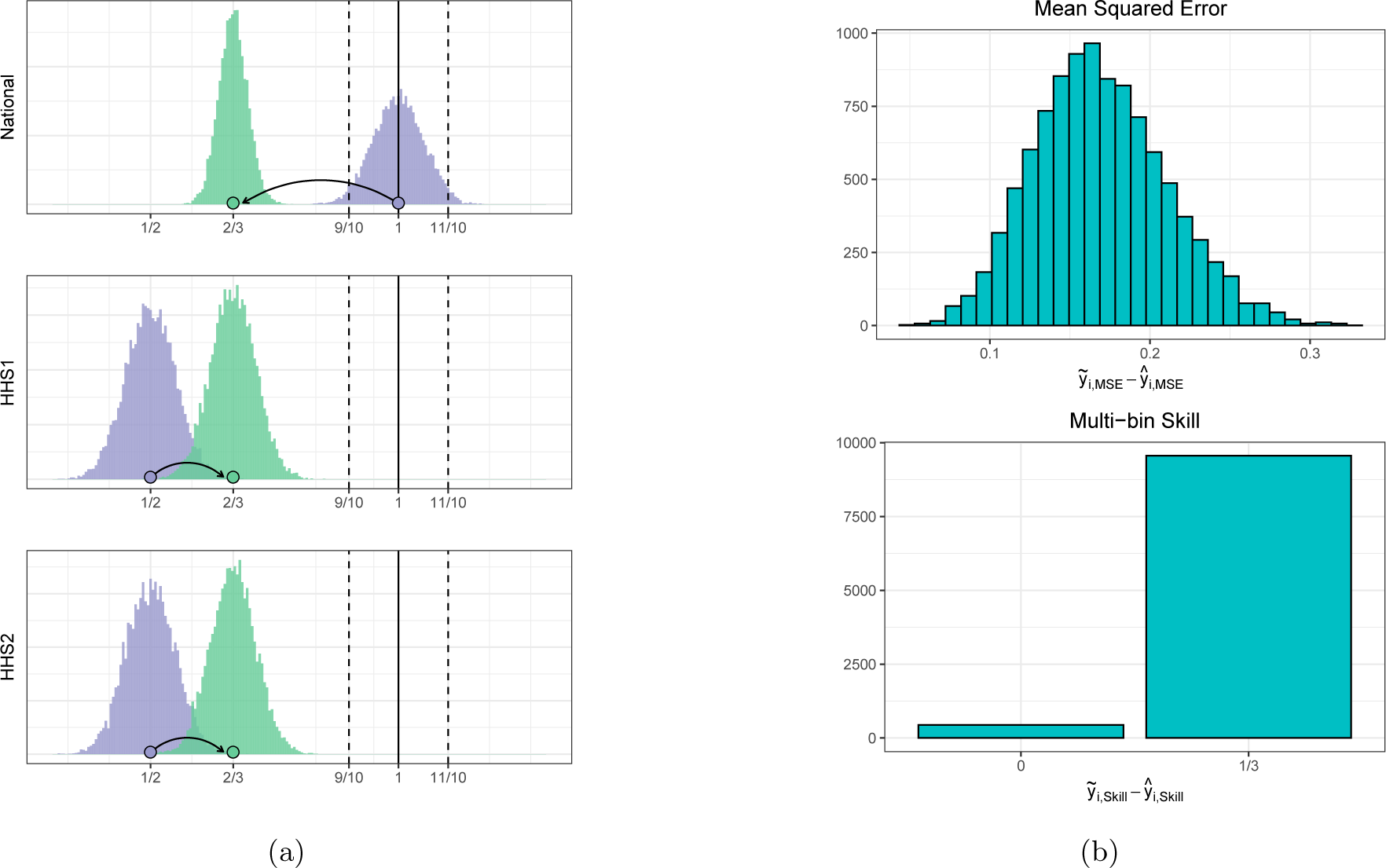
Graphical example of how mean squared error (MSE) can decrease while skill gets worse for two region example. **A:** Purple histograms represent the 10,000 realizations of 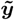, while green histograms are the corresponding ***ŷ***s. The purple and green points illustrate a particular example of the projection matrix forecasting process. The solid vertical lines denote the true value for each region, while the vertical dashed lines denote the correct regions for the skill. **B:** Top panel shows distribution of MSE for 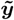 minus corresponding MSE for ***ŷ***. MSE for 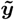 is greater than the MSE for ***ŷ*** for all realizations. **B:** Bottom panel shows multi-bin skill score for 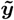 minus skill score for ***ŷ***. The incoherent 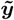 forecasts are better or equal to the skill for the coherent forecasts for all iterations.

## 2 Methods

In order to investigate the question posed in Section 1.4 we developed two methods to sample from probabilistically coherent forecast densities. To begin, we define our collection of original forecast densities, drawn independently for each region:

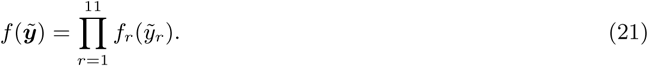

Previous approaches have factored 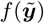 into a bottom-up density, where

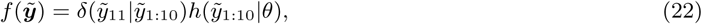

where *δ* is a Dirac delta density centered at 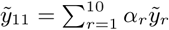, *θ* is a parameter(s) estimated from training data, and *h* is a joint density over all 10 regions [18].

The bottom-up model of Equation 22, while probabilistically coherent, lacks robustness in two key ways. First, it requires historical training data to estimate *θ*. This is not always possible, particularly in emerging epidemic settings. Second, the bottom-up approach ignores information encoded in the original forecast density for the national region 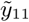. We instead develop methods that take draws from 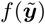 of Equation 21 and produce draws for a probabilistically coherent *f*^∗^(***ŷ***). These methods address both of the shortcomings of the bottom-up density of Equation 22: they do not discard national scale forecasts and they do not require training data.

In what follows we consider two approaches for sampling from the probabilistic coherent density, *f*^∗^(***ŷ***). We first consider the scenario where errors in the forecast distributions are assumed to be uncorrelated across regions. This allows us to sample from the original forecast distribution 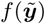 and apply a point-wise coherence projection matrix to each independently drawn sample 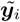. In the second approach, we assume the geographical units are positively correlated. That is, the forecasting model is more likely to underestimate or overestimate locations simultaneously rather than having the errors be independent of one another. This correlation structure reflects our knowledge that during an epidemic, forecast models that tend to under predict wILI at the regional level, will also do so at the national level.

### 2.1 Unordered Ordinary Least Squares

Our first approach requires samples from the independent forecast distributions, defined as follows:

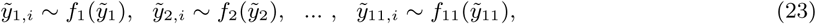

where *i* indexes samples and where we have dropped the season, epiweek, and model index for simplicity. We then apply the projection matrix to the column vector

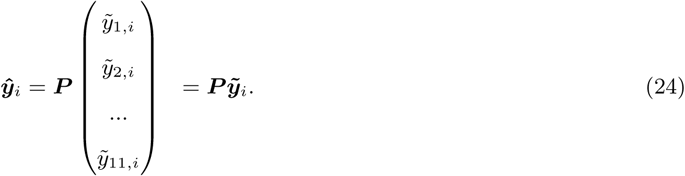

This approach is graphically demonstrated in Figure 4. Using the projection matrix on each sample guarantees that the resulting empirical probability mass function satisfies the probabilistic coherence definition of Equation 16. Algorithm 1 outlines how to produce samples from *f*^∗^(***ŷ***). In practice, the CDC submission files specify probability distributions as binned probability mass functions. Independent forecasts 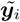 are sampled from these binned probability mass functions.

**Figure 4:**
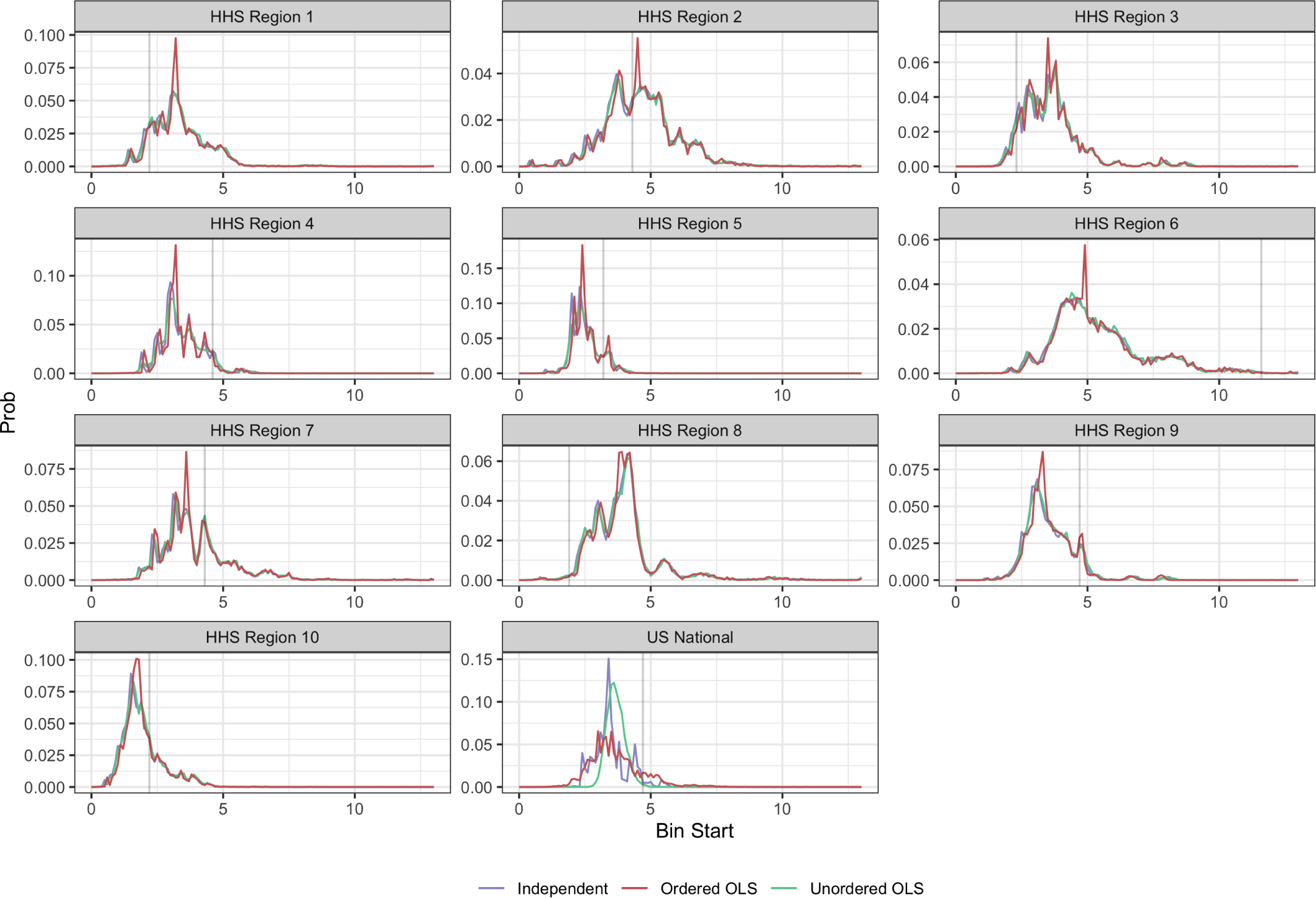
Real data example of model predictive densities for the 1 week ahead target on epiweek 201901 for the 2018/2019 season across all 11 regions. Purple lines show samples from the independent densities. The green lines represents a coherent transformation using the unordered ordinary least squares projection matrix on the purple samples. The red lines represents a coherent transformation using the ordered ordinary least squares projection matrix on the purple samples. Notice how the regional samples do not change much under the coherence constraint, but the national forecasts noticeably change. True wILI for epiweek 201902 shown in black for each region.

#### Algorithm 1

Unordered OLS sampling from probabilistically coherent joint distribution given a collection of marginal distributions.

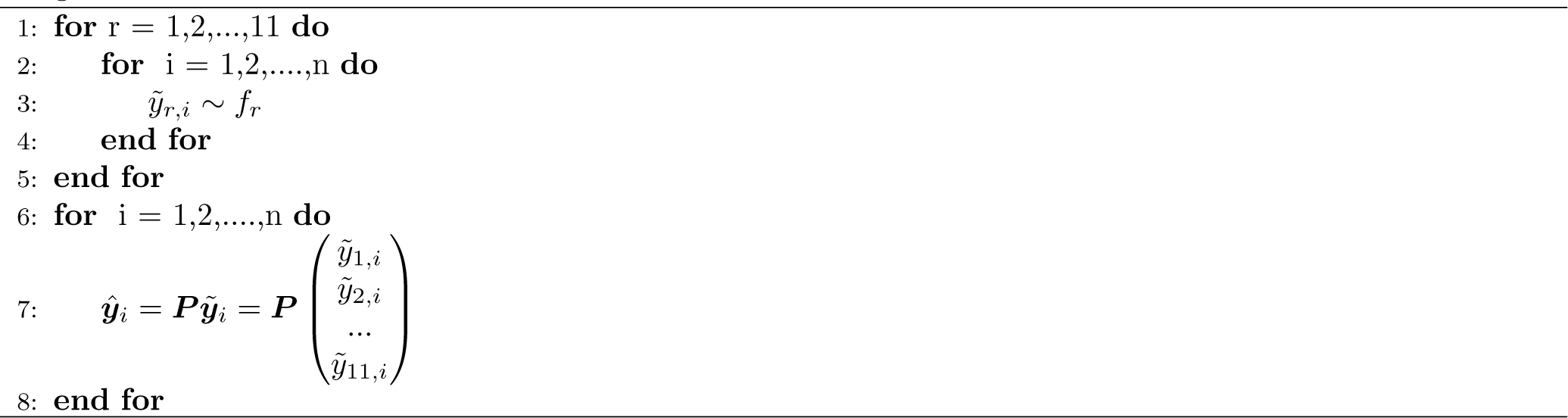

### 2.2 Ordered Ordinary Least Squares

The unordered OLS approach assumes no correlation between the error structures regionally and nationally. That is, each sample is generated independently. In the second approach we induce a correlation structure between the forecast errors. We begin with the same set of samples as defined in Equation 23. However, before applying the projection matrix we first compute the order statistics. We then apply the projection matrix to the column vector to the aligned order statistics:

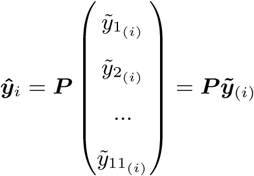

where 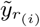 is the *i*^*th*^ and order statistic for the empirical distribution 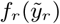 for region *r*.

Both the ordered and unordered OLS approaches lead to empirical distributions that are probabilistically coherent, however, the ordered OLS approach induces a correlation structure where low regional wILI forecasts are tied to low national wILI forecasts and vice versa: similar to the Schaake Shuffle [24]. In practice, the ordered OLS algorithm amounts to first sorting the samples drawn independently at each region and then applying the projection matrix to the sorted samples as outlined in Algorithm 2.

#### Algorithm 2

Ordered OLS sampling from probabilistically coherent joint distribution given a collection of marginal distributions.

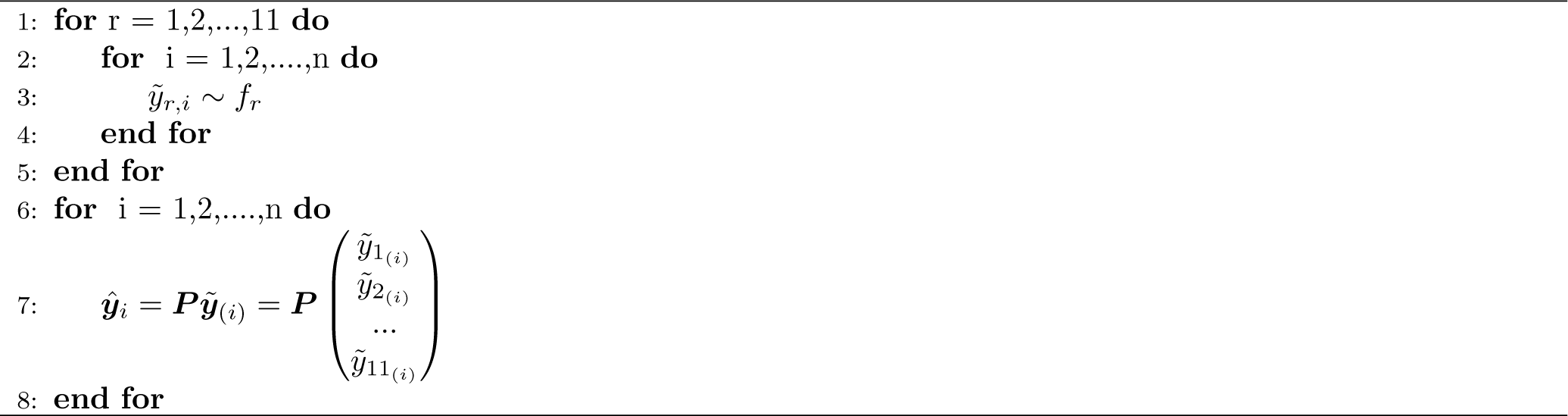

### 2.3 Experimental Setup

In order to examine the effects of the unordered and ordered OLS approaches on forecast skill, we use submission files for the 2016/2017, 2017/2018, and 2018/2019 seasons that have been submitted to the CDC and are uploaded to the central repository [17]. For each evaluation season, we obtain a list of all models submitted, and evaluate both approaches across epiweeks 44-17. Any model that did not have a complete set of submission files for all 1-4 week ahead targets, all epiweeks, and all regions was discarded. The sample sizes for the evaluation are included in Table 1, where we define an evaluation point as a unique region, season, model, epiweek, and target combination. As we can see from Table 1, the sample sizes are quite substantial when aggregating over evaluation points. We use the 2010 U.S. Census weights across all seasons as an estimate of *α*_*r*_ for each region [22]. The correlation between the weighted combination of the regionally reported wILI by the CDC and the nationally reported wILI is > .99. Using the 2010 U.S. Census weights is a reasonable approximation to the weights used by the CDC.

**Table 1:** Experimental setup for evaluating probabilistic coherence approaches. An evaluation point is defined as a unique region, season, epiweek, target, and model combination.

| Season | Number of Models | Number of Evaluation Points |
| --- | --- | --- |
| 2016/2017 | 24 | 27,456 |
| 2017/2018 | 24 | 27,456 |
| 2018/2019 | 35 | 40,040 |

## 3 Results

In what follows we consider the combination of a model and a season as the fundamental unit of analysis on which to base our conclusions. As a forecaster, the main question under consideration is whether applying forecast coherence will improve the average forecast skill of a given model in an upcoming season. As we can see from Figure 5 both the unordered and ordered OLS methods improved short-term forecasting skill for the majority of model/seasons, averaged over short-term targets, regions, and epiweeks. The ordered OLS method saw an improvement in two-thirds of the models submitted. However, the ordered OLS method showed improvement in forecast skill in 90% of models submitted.

**Figure 5:**
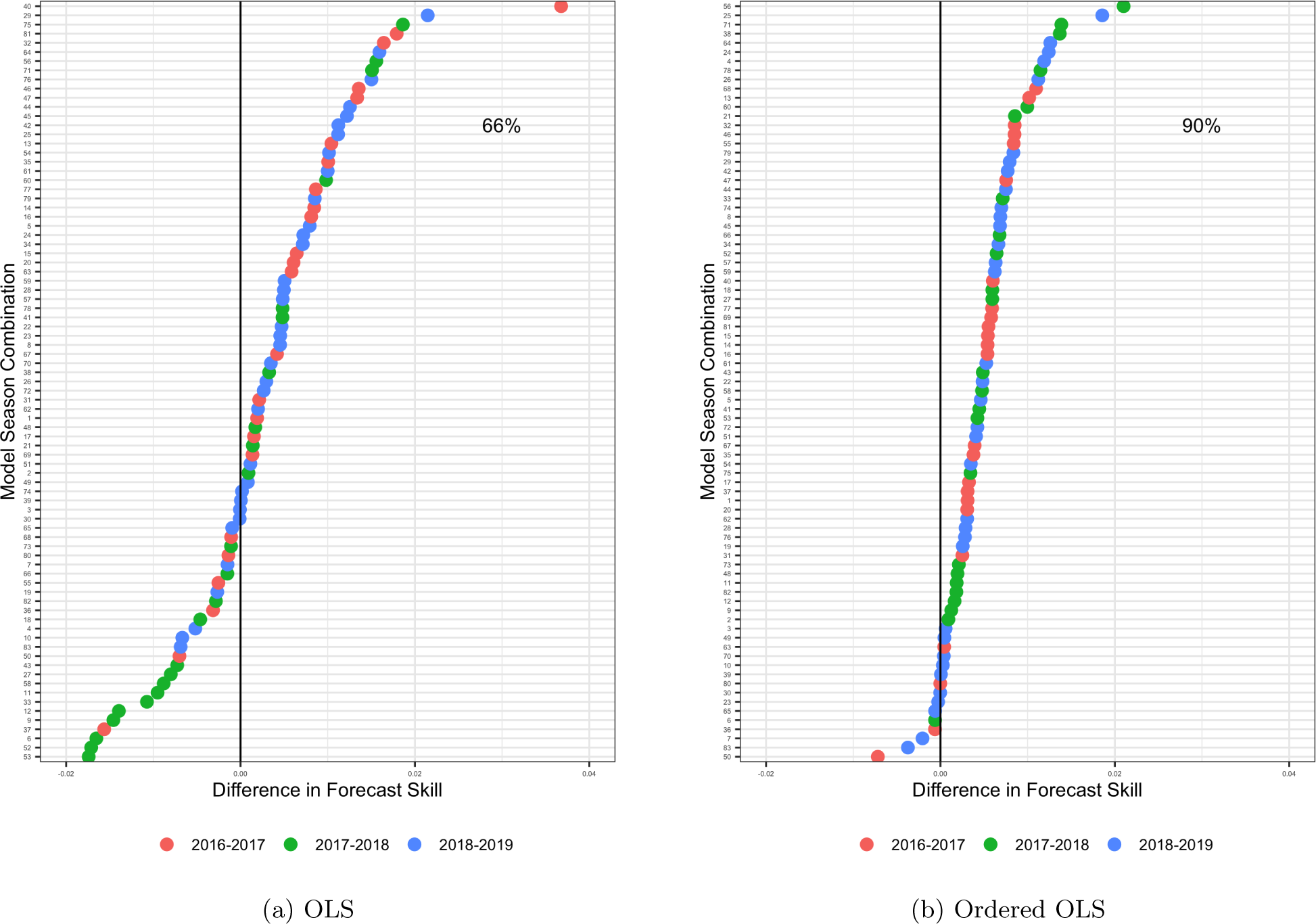
Difference between forecast skill of projection method and forecast skill of independent forecasts averaged over all targets (1-4 week ahead) and all test seasons (2016/2017,2017/2018,2018/2019) broken down by model. Notice that under the OLS method there is only a 66% chance of being better than the independent forecasts. he ordered OLS method, however, shows an even greater chance of improvement over the independent forecasts, with 90% of models performing better than their independent forecast counterparts..

The results are consistent across targets (see Figure 6 A), where a majority of model/season forecast skill improves over the forecast skill of the independent forecasts. Interestingly, forecast horizon has differing effects on the two methods, with ordered OLS improving as forecast horizon increase and unordered OLS improving as forecast horizon decreases. In particular, out of the 83 unique model/season combinations, the ordered OLS method improved over 75% of them for each target, showing that ordered OLS can consistently improve forecast skill across targets.

**Figure 6:**
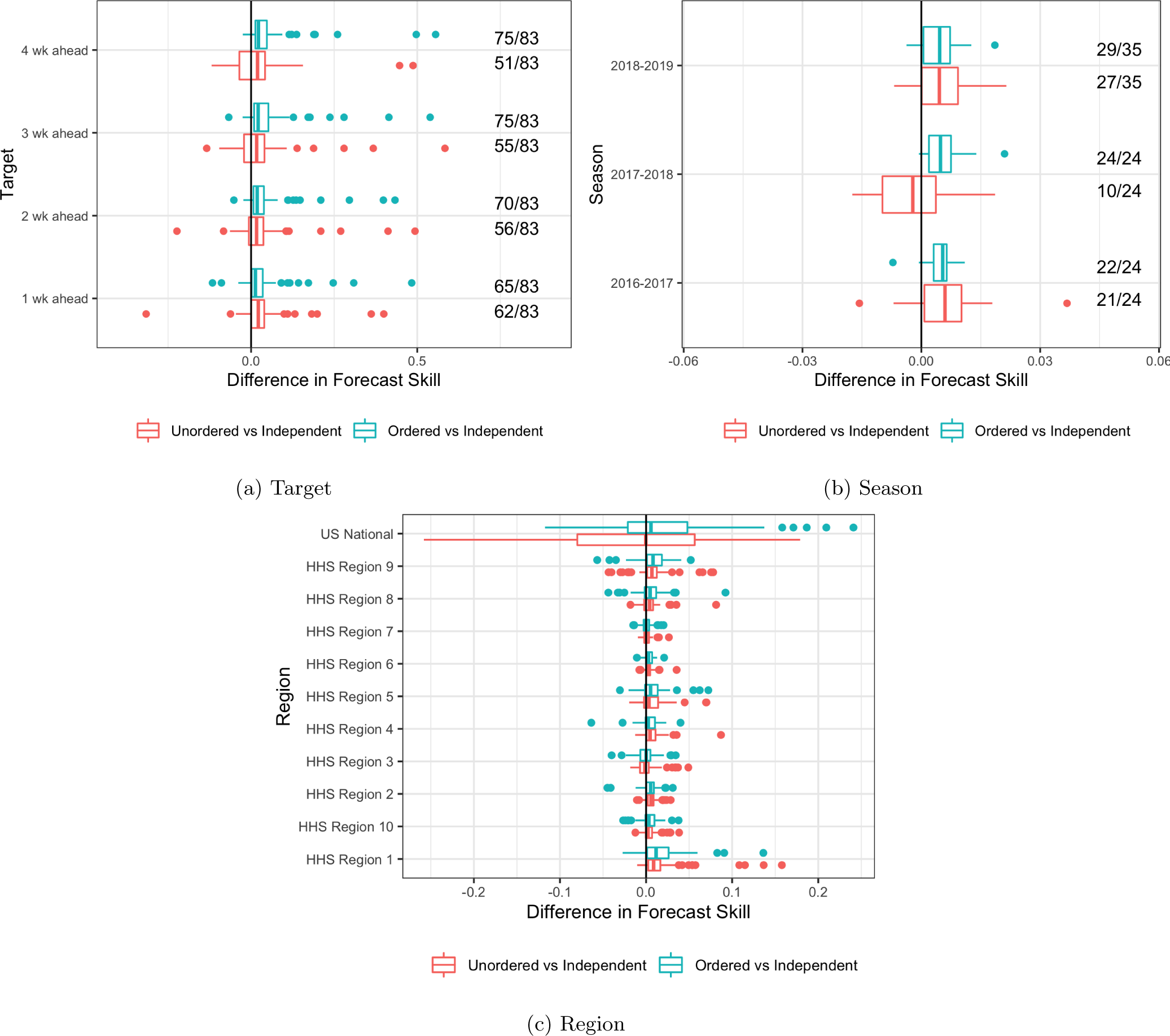
**A**: Difference between forecast skill of projection method and forecast skill of independent forecasts averaged over all regions and epiweeks broken down by target. Notice that only the ordered OLS method is significantly to the right of the zero line. Numbers represent the number of models that improved relative to the independent forecast skills. **B**: Difference between forecast skill of projection method and forecast skill of independent forecasts averaged over all regions, epiweeks, and targets (1-4 week ahead) broken down by season. Notice that the 2017/2018 season, which was the hardest of the three seasons to predict, saw the largest increase in forecast skill and the largest relative difference in the ordered OLS versus the unordered OLS. Numbers represent the number of models that improved relative to the independent forecast skills. **C**: Difference between forecast skill of projection method and forecast skill of independent forecasts averaged over epiweeks and targets (1-4 week ahead) broken down by region. Here we see no systematic improvement. However, we do not expect to see one since forecast coherence does not necessarily improve the forecast skill of every region, but rather the regions in aggregate. When breaking down the results by region, the geometric mean obscures any average benefit across region.

We also break down the results by season (Figure 6 B). Here we see that both methods improve over the independent forecasts when stratified by seasons. Additionally, the ordered OLS method showed the greatest improvement in the season that was hardest to predict out of the three: the 2017/2018 season [12]. In fact, the top skills obtained by all participants in the FluSight challenge during the test seasons were .45, .33, and .44 respectively, representing a noticeable drop in forecast skill for the 2017/2018 season. The ordered OLS method was able to improve all 24 models submitted in the 2017/2018 season, while the unordered OLS method only improved 10 of the 24. This suggests that the benefits of coherence are most clear in seasons that are difficult to predict, a desirable property.

Though the ordered OLS method shows consistent improvement in short-term forecasting skill across model/seasons (74/83) and when broken down by target (Figure 6 A) and season (Figure 6 B), consistent improvement in forecast skill does not hold when broken down by region as seen in Figure 6 C. As demonstrated in Section 1.2, there is no guarantee that in the point prediction setting the MSE of each region will improve, but rather the average MSE across regions will improve.. This property seems to translate to the probabilistic forecasting realm as well. There is no systematic improvement in Figure 6 C. In fact, the breakdown obscures any improvement at all, a consequence of using forecast skill as the primary metric, which is a geometric mean of log score.

We next turn to the question of why the ordered OLS method performed better than the unordered OLS method. As noted in Section 2.2, the ordered OLS method assumes a correlation structure exists between the errors of the independent forecast densities. In particular, the ordered OLS method assumes that when the regional forecasts are low, the national forecast is similarly low and vice versa. When examining the observed correlation between the average error of the regional forecasts (averaged over all regions) and the error of the national forecasts in the evaluation seasons (Figure 7), we can clearly see positive correlation that the ordered OLS method is able to exploit. This helps explain the overall increase in performance seen with the ordered OLS method.

**Figure 7:**
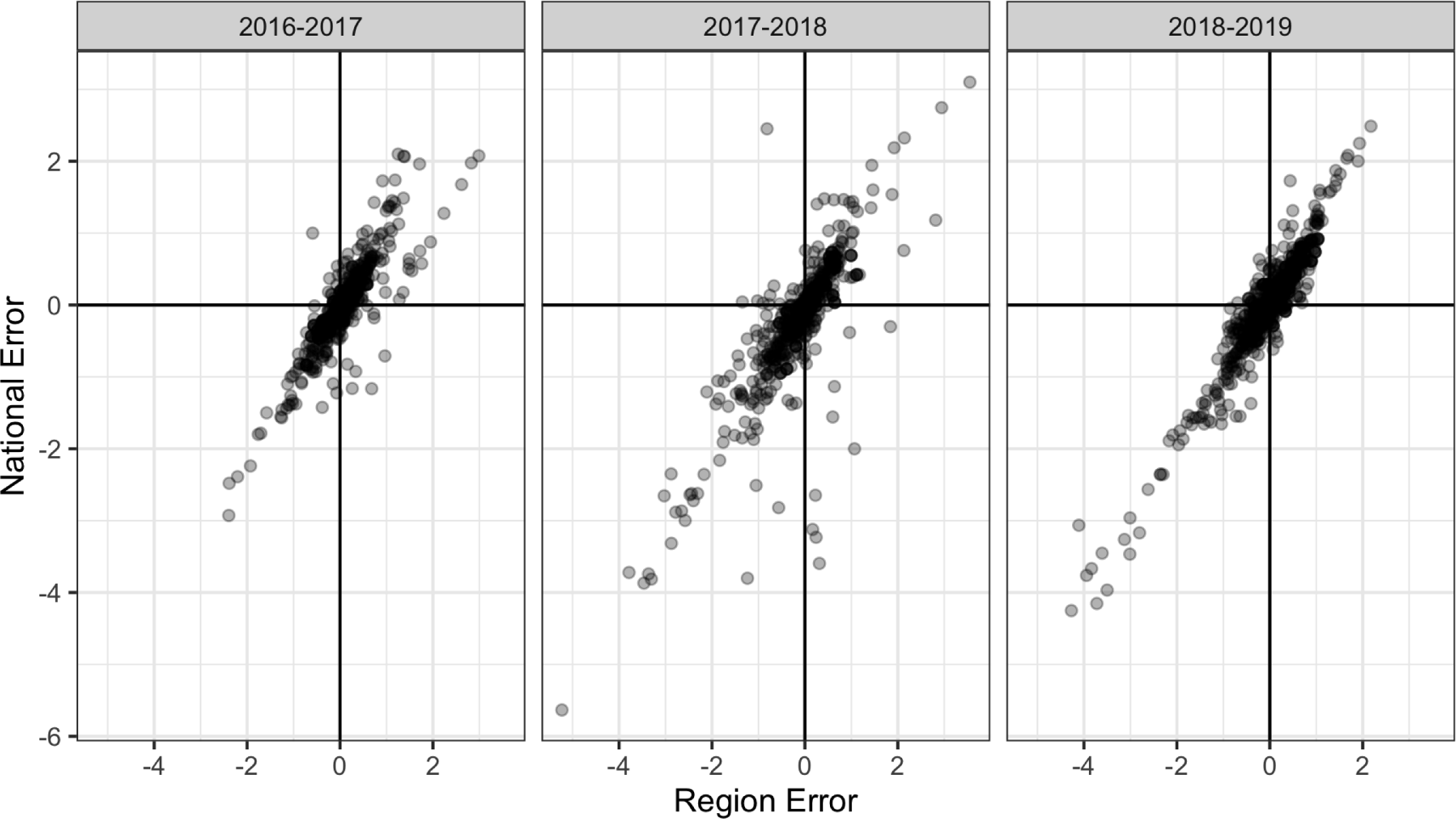
Average error of the original forecast distribution (averaged over region) against national error where each point corresponds to a model, season, epiweek combination averaged over all targets. Most of the points fall in both the 1st and 3rd quadrant, indicating that when the average regional error is high, the national error is likely to be high and vice versa. This explains why the ordered OLS method performs better than the unordered OLS method, since it is able to capture this correlation structure.

## 4 Discussion

Forecast coherence is a simple tool to improve forecast skill of short-term predictions in systems with hierarchical structures. In order to demonstrate this, we first defined probabilistic coherence, and showed that the results in the literature surrounding point forecast coherence do not naively transfer over to probabilistic forecasting. Guarantees for improvements in MSE do not directly transfer over to the CDC FluSight forecast skill metric of probabilistic forecast performance. However, by leveraging the definition of probabilistic coherence, we were able to generate coherent samples by first sampling from a collection of independent forecast distributions over all regions and projecting them onto a coherent subspace. This projection method is generic and allows for both correlated and uncorrelated projections. By exploiting the correlation structure of the true data generating process, we were able to improve average forecast skill when broken down by model, season and target.

In practice, the unordered and ordered OLS methods are very appealing due to the lack of training data required and the operational simplicity of operating on submitted forecast, without requiring adjustments to the model code itself. No knowledge of the process model used to generate forecasts is required, only the resulting predictive density. Even though the benefits in forecast skill are small in magnitude, there is little cost to implementing coherence in practice and, especially for the ordered OLS method, the frequency of improvement is high (90%).

Our experiments lead to the following conclusions:

- **Forecast coherence can benefit forecast skill, but the benefits are small**. Using the ordered OLS method, we can improve short-term forecast skill with high likelihood (90% of model/seasons) and when partitioned by targets (over 75% of all model/seasons improved for each target). We see a small but consistent improvement, with little to no cost in terms of parameter estimation and implementation difficulty. This makes the ordered OLS method a clear choice to use when submitting forecasts to the CDC FluSight challenge.
- **Careful consideration is required when accounting for correlation structure of the errors**. The ordered OLS method is able to account for positive correlation structure by applying the projection matrix the order statistics from the independent forecast distributions.
- **Coherence is most beneficial during a difficult season**. We saw that the forecast skill benefit for both methods was largest in the 2017/2018 season. In addition, the relative difference between the unordered OLS method and the ordered OLS method was also largest during this season, suggesting that exploiting the correlation structure matters most in difficult seasons [12]. This is intuitively true, since during an epidemic season both the regions and the national wILI is elevated beyond normal levels, so the forecasting models tend to under-predict at all levels of the hierarchy.

Although the above method is simple to implement and results in small but significant changes in forecast skill, there is still significant room for improvement. Recent work by Taieb et. al has explored copula based techniques to combine the independent forecast distributions from the regional level into a joint distribution with a specified covariance structure [18]. Wickramasuriya et. al have also explored various projections using a weighted least squares method [19]. This weight matrix also represents the correlation structure between the independent forecasts but can be estimated from historical forecast accuracy. It is clear that, unlike in the point forecast setting, exploring various correlation structures in the probabilistic setting has a drastic effect on the results. Given historical training data, once could estimate the error correlation specific to a given process model which could potentially lead to an even greater increase in forecast skill. However, in the absence of historical training data, forecasters can still leverage coherence to improve probabilistic forecast skill. In all, these results suggest that simple and fast methods can improve probabilistic forecasts of systems where the available data has a natural hierarchy. Using the example of seasonal influenza forecasts in the US, we show that enforcing coherence provides a high likelihood of improvement in forecast accuracy, and in general may provide opportunities for improvement in forecast accuracy in this and other real-world application settings.

## 5 Appendix

### Theorem 1.

*Let* **X**_*n×p*_ *be a matrix of full column rank*. *Assume **y** ∈ colspace(**X**)*, 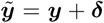 *where **δ*** ∈ ℝ^*n*^, and 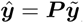, *where **P*** = ***X***(***X***^*T*^***X***)*−*1***X***^*T*^. Then, 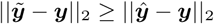.

*Proof*. Properties for the proof:

Property 1: Let ***X*** be an *n × p* matrix such that rank(***X***) = *p*.

Property 2: Define ***P*** = ***X***(***X***^*T*^***X***)^*−*1^***X***^*T*^. Then

Property 2a: ***P*** is idempotent (i.e., ***P*** = ***PP*** = ***P*** ^2^).

Property 2b: ***P*** is symmetric (i.e., ***P*** = ***P***^*T*^).

Property 2c: ***I − P*** is idempotent.

Property 2d: ***I − P*** is symmetric.

Property 3: If ***A*** is a symmetric matrix, then ***A*** = ***Q*Λ*Q***^*T*^ where ***Q*** is a matrix whose columns are equal to the eigenvectors of ***A*** and **Λ** is a diagonal matrix whose diagonal elements *λ* are the eigenvalues of ***A***.

Property 4: If ***A*** is idempotent, then its eigenvalues are either 0 or 1.

Property 5: Assume ***y*** ∈ colspace(***X***).

Property 6: 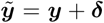, where ***δ*** ∈ ℝ^*n*^.

Property 7: 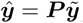

The proof proceeds as follows:

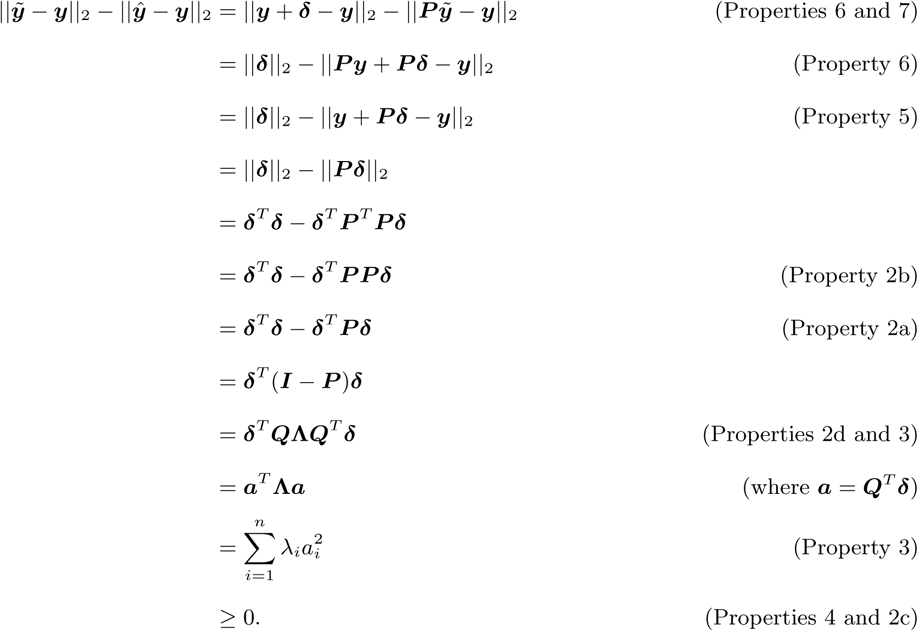

Thus, 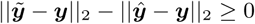 implies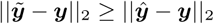.

## 6 Acknowledgements

This work was funded by the Department of Energy at Los Alamos National Laboratory under contract 89233218CNA000001 through the Laboratory-Directed Research and Development Program, specifically LANL LDRD grant 20190546ECR. The authors thank C.C. Essix for her encouragement and support of this work, as well as the helpful conversations with Dr. Brian Weaver. Approved for unlimited release under LA-UR-19-32598.

